# Botulinum Neurotoxin A1 Signaling in Pain Modulation within Human Sensory Neurons

**DOI:** 10.1101/2025.06.18.660472

**Authors:** Katherin A. Gabriel, Kali Hankerd, Paulino Barragan-Iglesias, Amy D. Brideau-Andersen, Lance Steward, Steve McGaraughty, Edwin Vazquez-Cintron, Theodore J. Price

## Abstract

Botulinum neurotoxin type A1 (BoNT/A1) is an effective treatment for chronic migraine, but its direct mechanism of action on human sensory neurons has not been fully elucidated. While rodent studies on dorsal root ganglion (DRG) and trigeminal ganglion (TG) show that BoNT/A1 inhibits neurotransmission, including calcitonin gene-related peptide (CGRP) release, by cleaving SNAP-25, only one previous study has assessed its effect on human DRG neurons. The objective of this study was to understand the mechanism of action of BoNT/A1 in cultured human sensory neurons and assess, using RNA sequencing, the transcriptomic consequences of BoNT/A1 treatment. Using DRGs obtained from organ donors the expression of key targets, including SNAP25, SV2C, & CALCA, was validated by mining existing transcriptomic datasets as well as immunohistochemistry. Cultured dissociated human DRG neurons treated with BoNT/A1 were used to examine cleavage of SNAP25, release of CGRP and transcriptomic changes after BoNT/A1 treatment. SV2C was found to be widely expressed in human DRG neurons in a pattern that completely overlapped with CGRP expression. Consistent with this finding, BoNT/A1 disrupted SNARE protein complexes in human DRG neurons as demonstrated by SNAP-25 cleavage in most somatosensory neurons and a reduction in capsaicin-evoked CGRP release, indicating impaired vesicle fusion. Moreover, Bulk RNA sequencing experiments revealed downregulated expression of a large subset of genes responsible for neurotransmitter and neuropeptide release from neurons suggesting a novel mechanism through which BoNT/A regulates neurotransmission. These results provide new insight into the molecular mechanisms by which BoNT/A may exert its pain-relieving effects in humans.

## Introduction

Botulinum toxins (BoNTs) are neurotoxins that were originally described as potent inhibitors of neuromuscular function given that when injected intramuscularly, the release of acetylcholine is blocked at the neuromuscular junction leading to long lasting, reversible, muscle paralysis (Pantano & Montecucco 2014; SCOTT *et al*. 1973). Since their original discovery, BoNTs have been widely used as pharmaceutical agents.

OnabotutlinumtoxinA (onabotA., BOTOX®), is FDA-approved to treat neuronal hyperexcitability in various disorders, including motor conditions such as spasticity, autonomic disorders like overactive bladder, and sensory disorders like chronic migraine (Brin & Blitzer 2023) in addition to several upper facial aesthetic concerns. OnabotA is currently used to reduce headache frequency and improve patient quality of life in patients with chronic migraine (Aurora *et al*. 2010; Aurora *et al*. 2011; Diener *et al*. 2010; Dodick *et al*. 2010; Lipton *et al*. 2011; Silberstein *et al*. 2013). The clinical efficacy of onabotA in migraine is at least attributed to a direct effect on nociceptors as multiple studies have demonstrated that onabotA treatment decreases pain-like responses in animal models and also decreased CGRP release from rodent TG neurons (Moore *et al*. 2023; Olbrich *et al*. 2017; Blanshan *et al*. 2019; Navratilova *et al*. 2022).

BoNTs block the release of acetylcholine, glutamate, and neuropeptides like CGRP via cleavage of soluble N-ethylmaleimide-sensitive factor attachment protein receptor (SNARE) protein complexes in peripheral and central neurons (Pellett *et al*. 2015). Synaptosomal-associated protein 25 (SNAP-25), an important protein part of the SNARE complex, is the target of BoNT/A1, and measuring the cleavage of this protein is a marker for BoNT/A1 action within a neuron. SV2 proteins A-C are the cell entry proteins for BoNT/A1 uptake. Sv2c is the primary protein receptor for BoNT/A1 entry (Dong *et al*. 2006; Mahrhold *et al*. 2006). A previous study in human DRG neurons, and in nerve samples from human clinical samples, reported extensive Sv2a immunoreactivity that appeared to increase with inflammation and/or nerve injury (Yiangou *et al*. 2011). However, recent RNA sequencing studies in human DRGs and TG suggest that SV2C is far more highly expressed compared to SV2A and SV2B, and the SV2C gene is expressed by all subtypes of somatosensory neurons in human DRG and TG (Tavares-Ferreira *et al*. 2022b; Bhuiyan *et al*. 2024; Yang *et al*. 2022). One of the goals of our study was to validate SV2C expression in the human DRG.

Despite the valuable insights that pre-clinical rodent models have provided into the mechanism of action of BoNT/A1 for pain and neuroinflammation (Cui *et al*. 2004; Meng *et al*. 2009; Carmichael *et al*. 2010; Gazerani *et al*. 2010; Yiangou *et al*. 2011; Coelho *et al*. 2014; Edvinsson *et al*. 2015; Coelho *et al*. 2016; Joussain *et al*. 2019; Melo-Carrillo *et al*. 2019; Melo-Carrillo *et al*. 2021; Belinskaia *et al*. 2022; Munoz-Lora *et al*. 2022) there remain many unanswered questions for the action of this biologic on human sensory neurons. The only previous study that examined human DRG neurons, found that a BoNT/A, from an unspecified source, modulated TRPV1-mediated Ca^2+^ signaling and neurite outgrowth (Yiangou et al, 2011; J Pain Res). Here, we show that the BoNT/A1 entry receptor Sv2 is highly expressed in all human DRG neurons.

Furthermore, BoNT/A1 treatment led to broad cleavage of SNAP25 across these cultured human DRG neurons, and this treatment also reduced baseline and capsaicin-evoked CGRP release.

Finally, we demonstrate a profound transcriptional change in human DRG neurons treated with BoNT/A1 suggesting a previously unknown mode of action of this toxin in regulating sensory neuron transmission and excitability.

## Material & Methods

### Bioinformatic analysis

We analyzed approximated whole-cell transcriptomes for single neurons as described previously (Tavares-Ferreira *et al*. 2022b) and DRGs from donors 1 to 8 were used in the analysis shown here. We also analyzed data from a recently published harmonized atlas of DRG and TG neuron transcriptomes across multiple species, and including human data from multiple studies (Bhuiyan *et al*. 2024).

### Human DRG recovery

All human tissue procurement procedures were approved by the Institutional Review Board at the University of Texas at Dallas. Human DRGs were surgically extracted using a ventral approach (Valtcheva *et al*. 2016) from organ donors within 4 hours of cross-clamp and placed immediately on artificial cerebrospinal fluid (aCSF). All tissues were recovered in the Dallas area via a collaboration with the Southwest Transplant Alliance as described in a previously published protocol (Shiers *et al*. 2024). The demographic information for the donors used in culture can be found in **Table 1**

**Table 1.** Demographic information.

| **Donor ID** | **Age** | **Sex** | **COD** |
| --- | --- | --- | --- |
| UTD-DN0302 | 43 | M | MVA |
| UTD-DN0303 | 18 | M | GSW |
| UTD-DN0292 | 22 | M | Anoxia/OD |
| UTD-DN0318 | 20 | M | MVA |
| UTD-DN0267 | 36 | M | MVA |
| UTD-DN0269 | 29 | F | GSW |
| UTD-DN0273 | 23 | M | MVA |
| UTD-DN0346 | 42 | F | Anoxia |

\*COD: cause of death
\*OD: Overdose
\*MVA: motor vehicle accident
\*GSW: Gunshot wound

### Human DRG dissociation protocol

The recovered DRG was trimmed with No. 5 forceps (Fine Science Tools, Cat#11252-00) and Bonn scissors (Fine Science Tools, Cat# 14184-09) to expose the cluster of cell bodies. At the time of culturing, a final enzyme solution consisting of 2mg/mL Stemzyme I (Worthington Biochemical, Cat#LS004106) and 0.1 mg/mL DNAse I was prepared in HBSS (Thermo Scientific, Cat#14170161) and allowed to incubate in a water bath. A total of 5 mL of enzyme solution was used to dissociate the hDRG tissue. Ganglia were finely minced with scissors into smaller fragments less than 3 mm (about 0.12 in). The tissue fragments were transferred to the prewarmed enzyme solutions in 10 mL conical tubes and placed in a 37°C shaking water bath until the tissue completely dissociated resulting in a cloudy solution (6-18 hrs in total). The solution containing dissociated neurons was passed through a 100 μm sterile cell strainer (VWR, Cat# 21008-950) into a 50 mL conical tube. The cell suspension was then gently added to a 3 mL 10% Bovine Serum Albumin (Biopharm, Cat#71040)/HBSS gradient and spun down at 900*xg* for 5 min at RTP, 9 acceleration, 5 deceleration. The resulting pallet of cells at the bottom of the tube was resuspended in prewarmed and sterile filtered DRG media. 100ul of cell suspension was carefully plated onto each coverslip, and 500μL onto the 6 well plates at a density of at least 150 and 800 neurons respectively and allowed to adhere for 3 hours at 37°C and 5% CO2. The wells were flooded with the appropriate amount depending on the plate of prewarmed complete hDRG media. Media changes were performed every other day.

### Plating Media

BrainPhys® media (Stemcell technologies, cat. no. 05790), 1% penicillin/streptomycin (Thermo Fisher Scientific, Cat# 15070063), 1% GlutaMAX® (United States Biological, cat. no. 235242), 2% NeuroCult™ SM1 (Stemcell technologies, cat. no. 05711), 1% N-2 Supplement (Thermo Scientific, cat. no. 17502048), 2% HyClone™ Fetal Bovine Serum (ThermoFisher Scientific SH3008803IR), 1: 1000 FrdU, 10 ng/mL Human β nerve growth factor - (Cell Signaling Technology, Cat# 5221SC)

### Immunohistochemistry

The human DRGs were gradually embedded in OCT in a cryomold by adding small volumes of OCT over dry ice to avoid thawing. All tissues were cryostat sectioned at 20 μm onto SuperFrost Plus charged slides. Sections were only briefly thawed to adhere to the slide but were immediately returned to the −20°C cryostat chamber until completion of sectioning. The slides were then immediately utilized for histology. Slides were removed from the cryostat and immediately transferred to cold 10% formalin (4°C; pH 7.4) for 15 minutes. The tissues were then dehydrated in 50% ethanol (5 min), 70% ethanol (5 min), 100% ethanol (5 min), 100% ethanol (5 min) at room temperature. The slides were air dried briefly and then boundaries were drawn around each section using a hydrophobic pen (ImmEdge PAP pen, Vector Labs). When hydrophobic boundaries had dried, the slides were submerged in blocking buffer (10% Normal Goat Serum, 0.3% TritonX 100 in 1X Phosphate Buffer Saline (PBS)) for 1 hour at room temperature. Slides were then rinsed in 1x PBS, placed in a light-protected humidity-controlled tray and incubated in primary antibody SV2C (1:100 Novus Biologicals # H00022987-M01) and SNAP25 (1:100 Thermo Scientific # 701991), diluted in blocking buffer overnight at 4°C. The next day, slides were washed in 1X PBS and then incubated in their respective secondary antibody Alexa Fluor goat anti-mouse IgG 555 (1:2000; Invitrogen A-21449, lot 2186435), Alexa Fluor goat anti-rabbit IgG 488 (1:2000; Invitrogen A-11034, lot 2110499) with DAPI (1:5000; Cayman Chemical; Cat # 14285) diluted in blocking buffer for 1 hour at room temperature. Sections were then washed in 1X PBS, air dried and cover slipped with Prolong Gold Antifade reagent.

### RNA Scope in situ hybridization

RNAscope in situ hybridization multiplex version 1 was performed as instructed by Advanced Cell Diagnostics (ACD). Slides were removed from the cryostat and immediately transferred to cold (4°C) 10% formalin for 15 minutes. The tissues were then dehydrated in 50% ethanol (5 min), 70% ethanol (5 min) and 100% ethanol (10 min) at room temperature. The slides were air dried briefly and then boundaries were drawn around each section using a hydrophobic pen (ImmEdge PAP pen; Vector Labs). When hydrophobic boundaries had dried, protease IV reagent was added to each section until fully covered. The protease IV incubation period was optimized as recommended by ACD. Slides were washed briefly in 1X phosphate buffered saline (PBS, pH 7.4) at room temperature. Each slide was then placed in a prewarmed humidity control tray (ACD) containing dampened filter paper and a mixture of Channel and Channel 2 probes (50:1 dilution, as directed by ACD due to stock concentrations) was pipetted onto each section until fully submerged. This was performed one slide at a time to avoid liquid evaporation and section drying. The humidity control tray was placed in a HybEZ oven (ACD) for 2 hours at 40°C. The target of interest (SV2C) was set in Channel 1 in combination with CALCA (CGRP) probe in channel 2. Following probe incubation, the slides were washed two times in 1X RNAscope wash buffer and returned to the oven for 30 minutes after submersion in AMP-1 reagent. Washes and amplification were repeated using AMP-2, AMP-3 and AMP-4 reagents with a 15-min, 30-min, and 15-min incubation period, respectively. Slides were then washed two times in 0.1M phosphate buffer (PB, pH7.4). Slides were incubated in 1:5000 DAPI in 0.1M PB for 1 min before being washed, air dried, and cover slipped with Prolong Gold Antifade mounting medium. Slides were imaged at 10X magnification using the Olympus IX83 Fluorescent microscope.

### BoNT/A1 treatments

Cells were cultured as described above and allowed to be in culture for 3-4 days before any treatment was given. Non-clinical grade BoNT/A1 (AbbVie) was given at 2 different concentrations (30pM and 100pM) for the western blot and immunocytochemistry experiments. For the CGRP release and the RNA sequencing 100pM was used. The cells were exposed to the toxin 24 hours before any media changes happened.

### Immunocytochemistry

The treated cultures were immediately fixed with 10% formalin (ThermoFisher Scientific, Cat# 23-245684) for 15 min at room temperature and rinsed twice with 1X PBS. 1hour block was performed at room temperature with 10% NGS and 0.3% Triton X-100 in 1X PBS. The coverslips were then incubated overnight at 4°C with primary antibodies anti-peripherin (chicken 1:1000; EnCor, AB_2284443) SNAP25 (1:5000; Millipore Sigma, s9684) or Snap25-197 (1:500; Allergan, AB635). Coverslips were rinsed 3 times with 1X PBS for 15 min and incubated with secondary antibodies Alexa Fluor goat anti-chicken IgG 647 (1:2000; Invitrogen A-21449, lot 2186435), Alexa Fluor goat anti-rabbit IgG 488 (1:2000; Invitrogen A-11034, lot 2110499) and Alexa Fluor goat anti-mouse IgG 594 (1:2000; Invitrogen, A11032, lot 1985396) for 1h at room temperature. The coverslips were washed 3 times with 1X PBS and mounted with prolong gold onto uncharged glass slides and allowed to cure overnight. Slides were imaged at 10X magnification using an Olympus IX83 Fluorescent microscope.

### Western blotting and analysis

Human DRG protein lysates were collected in ice cold RIPA lysis buffer (PI89900SDS) supplemented with protease inhibitor cocktail (Millipore Sigma P8340-1ML) and phosphatase inhibitor cocktail 2 & 3 (Millipore Sigma P5726-1ML & P0044-1ML). The tissue lysate was centrifuged at 12,000× *g* for 10 min at 4 °C, then the supernatant was collected and used for determination of protein content with a BCA assay. Total protein (20-40 μg) from tissue supernatant was loaded into TGX precast gels (4–20% CriterionTM, BioRad) and transferred to a PVDF membrane (Millipore Sigma IPFL00010). After transfer, the membrane was blocked at room temperature for 1 h in blocking buffer (5% dry milk in Tris-buffered saline with tween 20 (TBST)). The membrane was incubated in diluted primary antibodies SNAP25 (1:10,000; Millipore Sigma, s9684) for 24h at 4 °C. The membrane was washed three times in TBST for 5 min each then incubated with peroxidase-conjugated secondary antibodies for one hour rocking at room temperature. The membrane was then washed in TBST for 5 min each and then incubated with BioRad clarity western ECL substrate and Millipore HRP substrate solutions and then imaged in a ChemiDoc MP from BioRad.

### CGRP release

Human DRG cells were cultured and plated on a 24 well plate (Fisher Scientific NC0397150). On day 3 post plating, they were treated with an inflammatory soup consisting of PGE2 at 0.05uM, Serotonin at 0.5uM, Histamine at 0.5uM, Bradykinin 0.5uM, Substance P 0.5uM for 24 hours, based on a previous study (Moore *et al*. 2023). Media was completely removed and toxin (100pM) or Vehicle was added to the media for another 24-hour treatment. The Toxin was then removed, and new media was added and left for 72 hours. The day of the stimulation experiment, the media is removed and warmed at 37°C Tyrode’s Buffer (NaCl 119 mM, KCl 2.5 mM, MgCl2 2 mM, CaCl2 2 mM, HEPES 25 mM, Glucose 30 mM) was then added to the cells. They were then placed back into the incubator for one hour to equilibrate to the new buffer before the experiment began. Fresh Tyrode’s buffer with added thiorphan DL (16uM, Santa Cruz CAS 76721-89-6) was added to the vehicle groups, or Tyrode’s buffer with thiorphan and capsaicin (1uM; Millipore Sigma M2028) was added and left on for 30 minutes in the 37°C incubator. Media was then collected and analyzed for amounts of CGRP release levels using manufacturer recommendations for a human CGRP EIA kit (Cayman Chemicals 589101).

### Bulk RNA Sequencing

The RNA extraction was performed on cultured human DRG neurons using a Qiagen RNAeasy Plus universal Mini Kit. The RNA yield was quantified using a ThermoFisher Scientific Nanodrop system, and RNA quality was analyzed by an Advanced Analytical Technologies fragment analyzer. Stranded mRNA library preparation and sequencing were completed in house at the University of Texas at Dallas Genome Core, 100-bp, single-end sequencing of RNA-seq libraries was performed on the Illumina Hi-Seq sequencing platform. Raw sequencing (FASTQ) files were processed with FastQC to examine Phred scores, per-base sequence, and duplication levels. Reads were soft-clipped (12 bases per read) to discard adapters and low-quality bases during alignment using STAR v2.7.6 with the GRCh38 human reference genome (Gencode release 38, primary assembly). Reads were also sorted with STAR and then deduplicated with sambamba v0.8.2. Genes in the ENCODE blacklist were filtered out prior to quantification using bedtools (v2.30.0), and StringTie v2.2.1 was used to obtain normalized count values as transcripts per million (TPM) for each gene of all samples. The feature counts function of the R package Rsubread (v2.14.2) was used to calculate raw counts for all genes. DESeq2 R package (1.42.1) was used to perform the differential expression analysis of the different treatment groups. Then, Gene Ontology (GO) and Kyoto Encyclopedia of Genes and Genomes (KEGG) enrichment analysis of DEGs were determined using the clusterProfler R package (4.10.1).

### Statistics and data reporting

For the quantitative analyses, the statistical significance of treatment effects was examined using one-way analysis of variance (ANOVA) followed by Tukey’s post hoc test; p-values <0.05 were considered statistically significant completed in GraphPad Prism 9 (GraphPad Software, San Diego CA). Differentially expressed genes (DEGs) between treatments were obtained using the DESeq2 R package (1.42.1), using an adjusted p-value (padj)

## Results

### *SV2C* expression pattern in hDRG neurons using single nuclei and spatial sequencing

Using the harmonized cross species cell atlas of DRG neurons (Bhuiyan *et al*. 2024), we examined the mRNA expression pattern of synaptic vesicle 2 protein (*SV2C*) and synaptosomal associated protein 25 (*SNAP25*). The *SNAP25* transcript was robustly expressed in every neuronal cluster as was *SV2C,* with an 83% of cells showing co-expression. Only 2% of human neurons did not express one of these genes (**Fig. 1a-c**). While *SV2A* and *SV2B* were also expressed in this dataset, their overlap with *SNAP25* was only 43% and 51%, respectively.

**Figure 1.**
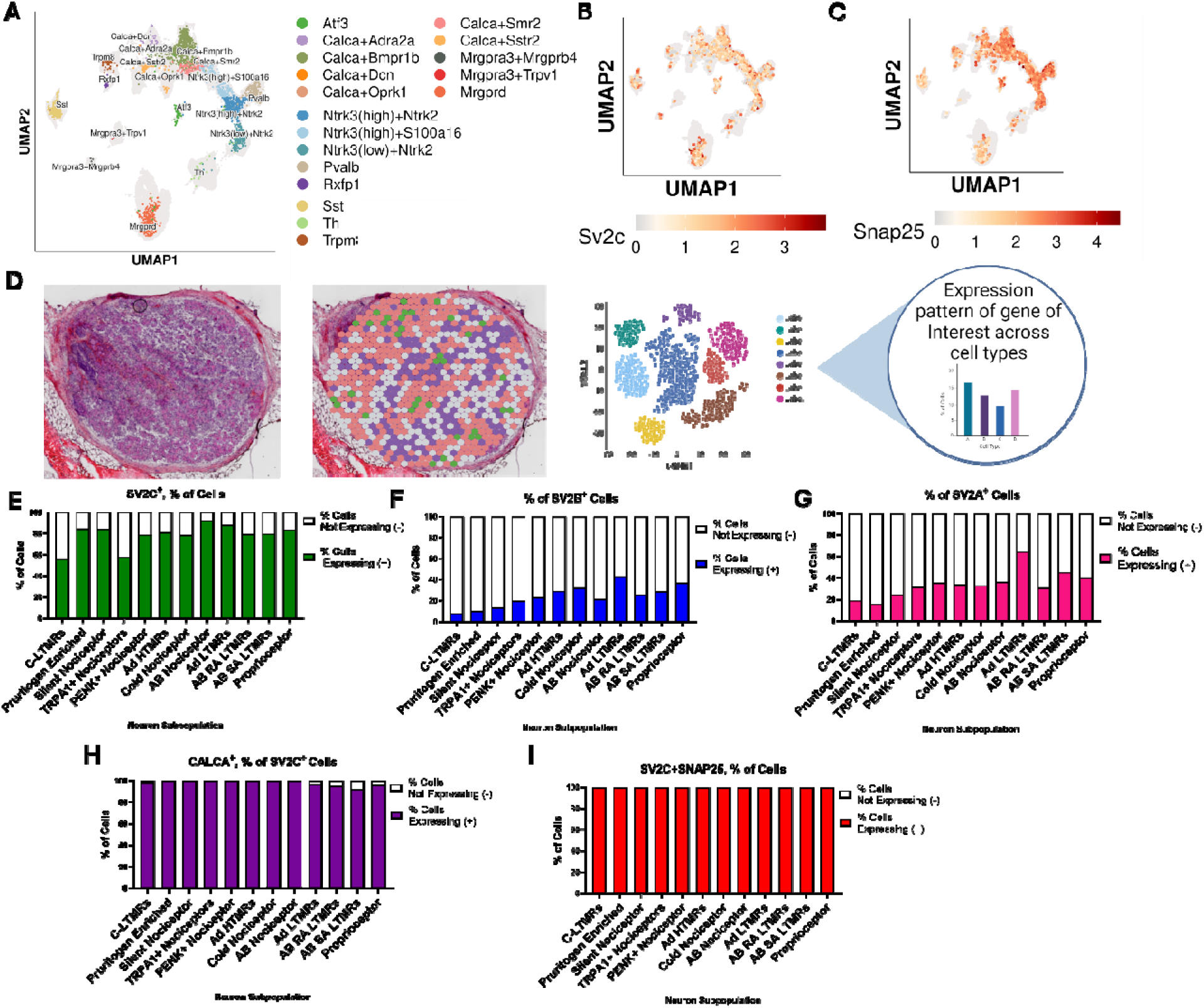
RNA sequencing expression of *SV2C, SNAP25* and *CALCA* in hDRG neurons. **A.** UMAP expression of all human clusters in the harmonized cross species cell atlas of DRG and the expression of **B**. *SV2C* and **C**. *SNAP25*. **D**. representative schematic of Visium spatial sequencing workflow. **E**. Percent of cells expressing *SV2C* **F.** *SV2B* and **G.** *SV2A* across 12 previously characterized neuronal subtypes. **H.** Percent of cells that are *CALCA* positive and express *SV2C*. **I.** Percent of cells that are *SV2C* positive and express *SNAP25*. Low threshold mechanoreceptors-LTMR, High threshold mechanoreceptors-HTMRs, Proenkephalin-PENK, Transient Receptor Potential Ankyrin 1-TRPA1.

Similarly, when reanalyzing a previously published 10X Visium dataset from human DRGs, it was found that *SV2C* was present in all 12 identified neuronal subpopulations with a high proportion of barcodes detecting *SV2C* in each sensory neuron subtype (**Fig. 1d-e**) (Tavares-Ferreira *et al*. 2022a). 55-90% of neuronal barcodes detected *SV2C* across neuronal populations with the lowest expression found in the C-LTMR subtype at 56% (**Fig 1e**). We see a much lower expression across all neuronal subtypes when looking at *SV2B* and *SV2A* transcripts (**Fig 1f&g**) When gene co-expression was analyzed, we see that 100% of the cells expressing *SNAP25* also express *SV2C*, and 98% of cells expressing the calcitonin gene related peptide (CGRP) transcript *CALCA* were also positive for *SV2C* (**Fig. 1h&i**). The quantity of reads for *SV2C* was higher in every single neuronal subtype (Tavares-Ferreira *et al*. 2022b) compared to *SV2B* and *SV2A*, demonstrating that Sv2c is the most robustly expressed entry receptor for BoNT/A in hDRG.

### Validation of SV2C and SNAP25 expression in hDRG tissue and culture

The expression of *SV2C* and SNAP25 was validated in hDRG tissue and in dissociated neuronal cultures from organ donor recoveries using immunohistochemistry, RNA scope and immunocytochemistry. Like the sequencing data, Sv2c and SNAP-25 proteins are highly expressed in the hDRGs and colocalized 100% (**Fig. 2a**). Using RNA scope, it was confirmed that *SV2C* transcript is expressed in 97% of the neurons, and from the *SV2C* positive cells, about 56% are *CALCA* positive as well, with all *CALCA*+ neurons also containing *SV2C* (**Fig. 2b-d**). The *CALCA* positive and *SV2C* positive cells had diameters ranging between 35µm and 85 µm, consistent with previous studies examining *CALCA* expression in human DRG (**Fig. 2e**) (Shiers *et al*. 2020a; Shiers *et al*. 2020b). We also validated the expression of SNAP-25 and Sv2c in eight-day *in vitro* dissociated neuronal cultures and compared the expression with the neuronal marker peripherin, a protein expressed in all neurons in the hDRG. The data demonstrates that most peripherin-positive hDRG neurons in culture express SNAP25 and Sv2c, allowing us to examine the effects of BoNT/A1 on SNAP-25 cleavage in hDRG cultures (**Fig. 2f & g**).

**Figure 2.**
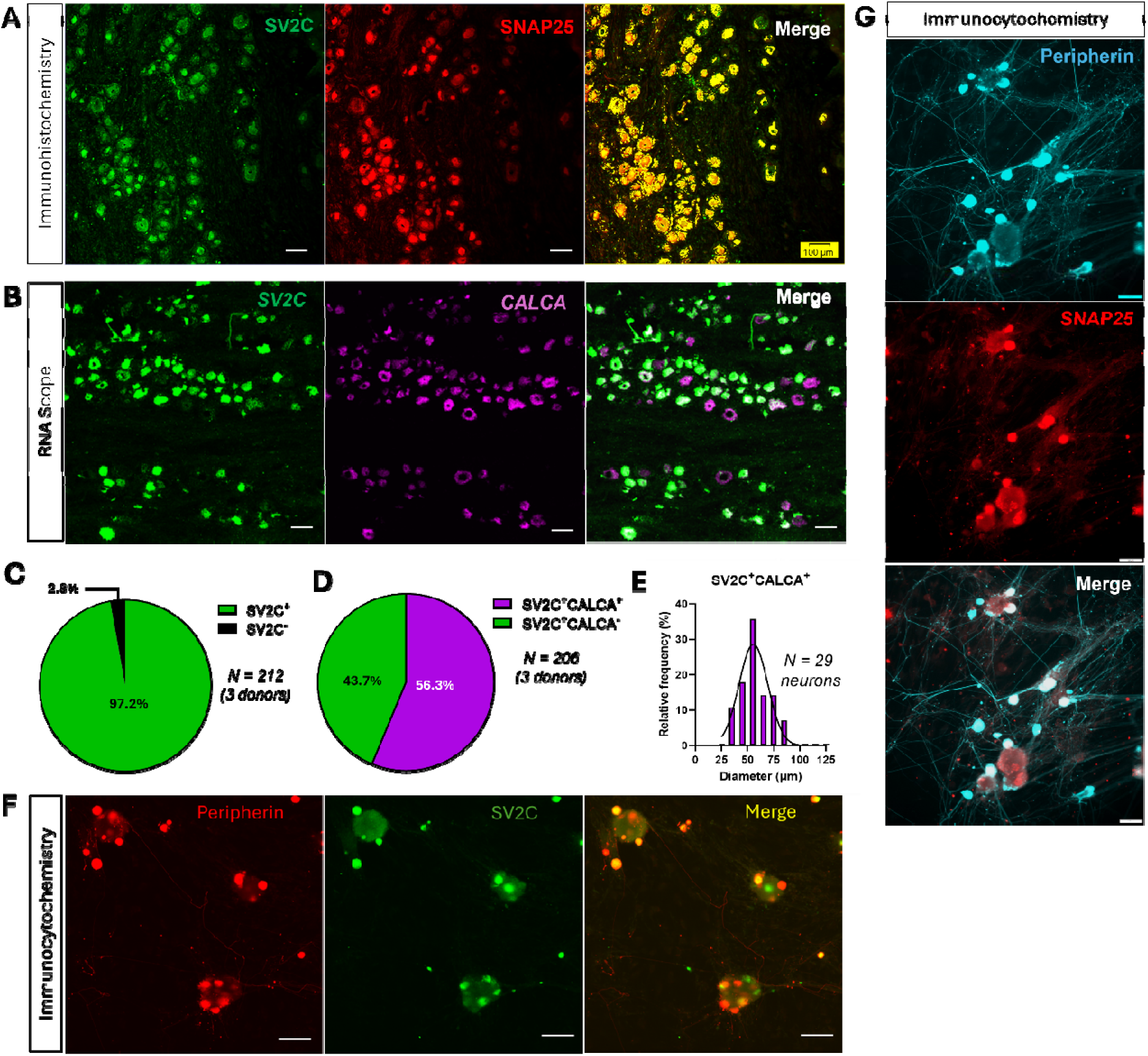
Protein and mRNA expression of *SV2C*, *SNAP25*, and *CALCA* in human DRG. **A**. representative 10x immunohistochemical images of SV2C (green), SNAP25 (red) and merge (yellow) in human DRG tissues. **B**. Representative 10x RNAscope in situ hybridization images of *SV2C* (green) and *CALCA* (purple) in hDRG tissue. **C**. Pie-chart representation of the percentage of neurons that are *SV2C* positive and negative across 3 individual donors in the RNAscope. **D**. Pie-chart representation of percentage of *SV2C* positive neurons that are either *CALCA* positive (Purple) or *CALCA* negative (Green) across 3 individual donors **E.** The frequency across the different diameters in *SV2C* positive and *CALCA* positive neurons. **F**. Representative 10x immunofluorescent images of Sv2c (green) and neuronal marker peripherin (red) in hDRG dissociated cultured neurons Representative **G.** 10x immunofluorescent images of SNAP25 (red) and neuronal marker peripherin (cyan) in hDRG dissociated cultured neurons. Scale bar=100 um

### BoNT/A1 treatment on hDRG cultures cleaves SNAP-25 in a dose dependent manner

Previous studies have shown that BoNT/A1 cleaves SNAP-25 in primary rodent trigeminal ganglion (TGs) cultures (Moore *et al*. 2023). Here, we used primary hDRG cultures to determine if this effect is recapitulated using BoNT/A1 on human neurons. At four-day *in vitro*, hDRG neurons were exposed to BoNT/A1 and SNAP-25 cleavage was measured using either immunohistochemistry or Western blot. Using an antibody that specifically detects the cleaved form of SNAP-25 (SNAP-25_197_), we found that SNAP-25197 only appears in the toxin treated cells compared to the vehicle (**Fig. 3a**). When comparing the co-expression of SNAP-25197 with the neuronal marker peripherin in the BoNT/A1 treated cells, nearly 40% of neurons in the 30pM treatment group showed SNAP-25197, and about 80% in the 100pM group showed SNAP-25197 (**Fig 3b**). Using an antibody that detects both the cleaved and the un-cleaved version of SNAP-25 in a Western blot, it was found that only the 100pM concentration of BoNT/A1 was sufficient to detect the -cleaved band via a western blot (**Fig. 3c&d**). This indicates that BoNT/A1 treatment cleaves SNAP-25 in hDRG dissociated neuronal cultures in a concentration-dependent manner.

**Figure 3.**
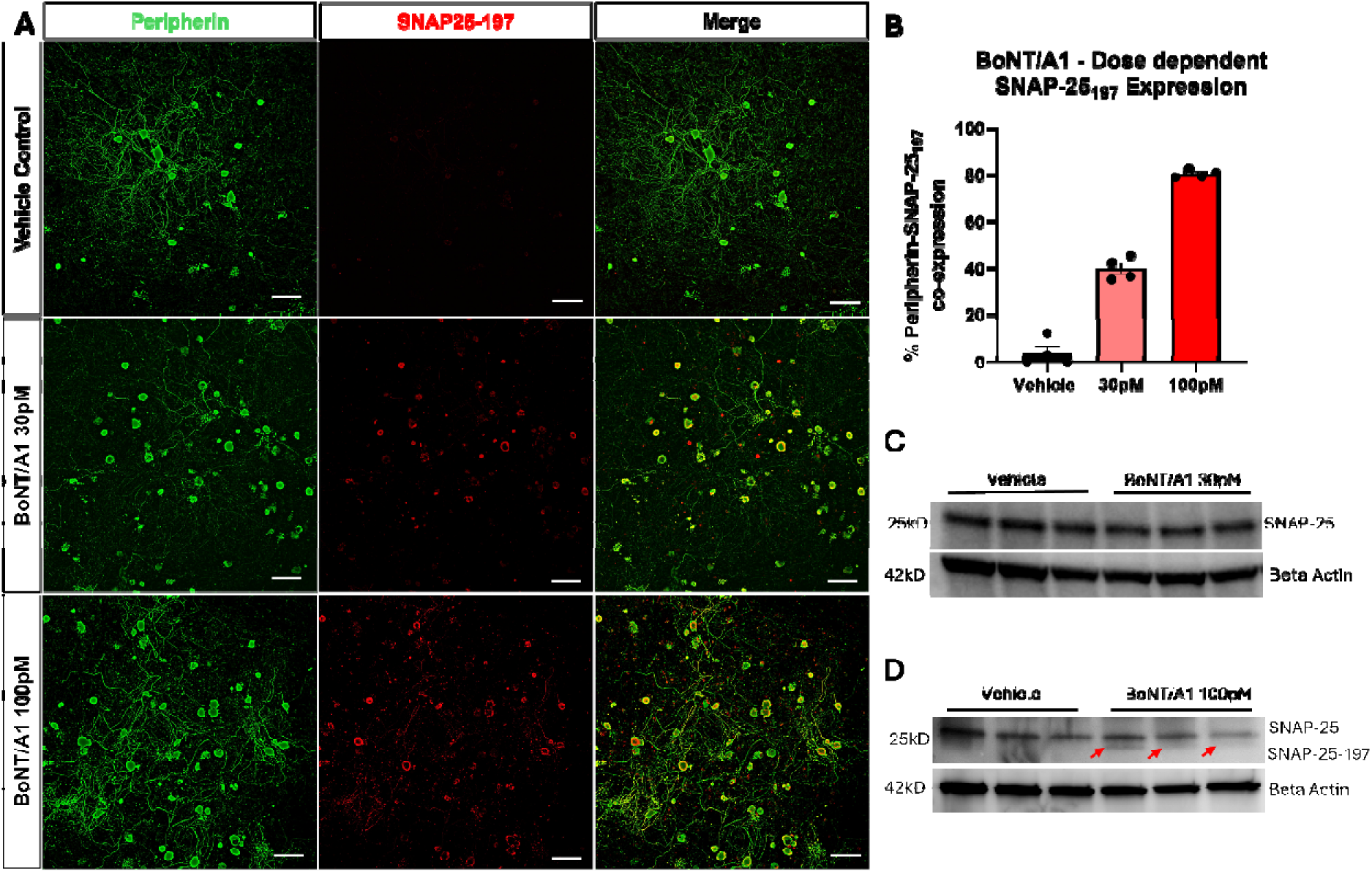
BoNT/A1 cleaves SNAP-25 in hDRG dissociated neurons in a concentration dependent manner. **A.** representative 10x images of dissociated hDRG neurons treated with vehicle or BoNT/A1 at 30pM or 100pM concentration. We used the neuronal marker peripherin (green), cleaved SNAP25 (red) and the merge of the two channels**. B.** quantification of the % of co-expression between cleaved SNAP-25_197_ and peripherin positive neurons across all three treatment groups. Each data point represents the average of individual replicates with at least 40 neurons counted from 3 different donors. Data is shown as means ± SEM. **C-D.** Representative western blot with antibody that detects both cleaved and un-cleaved SNAP-25 plus loading control Beta-actin comparing vehicle to 30pM and 100pM BoNT/A1 treatment. Scale bar=100um.

Due to the greater effect of 100pM BoNT/A1, we used this concentration of BoNT/A1 in subsequent experiments.

### Bulk RNA sequencing on BoNT/A1 treated hDRG cultures

Next, we wanted to determine what transcriptional changes occur between naïve cultures and cultures treated with 100pM BoNT/A1 through bulk RNA sequencing. We treated four-day *in vitro* hDRG cultures with BoNT/A1 or Vehicle for 24 hours and then waited 72 hours before extracting the RNA and then constructed libraries and sequenced them using the Illumina platform. The reads obtained were mapped to the reference human genome sequences. DESeq2 analysis was used to determine differentially expressed genes (DEGs) between the two groups. The principal component analysis (PCA) shows the similarity between biological replicates and variability between sample groups (**Fig. 4a**). We found a total of 273 DEGs, 15 upregulated and 258 downregulated (**Fig. 4b-d**). Among the downregulated genes in the BoNT/A1 treated cells, we found *SV2C, VAMP, SNAP25, SYT1* and *SYT2* which are all involved in synaptic vesicle trafficking and neurotransmitter release (adj. p-value <0.05). Similarly, *CACNA1B, KCNA1,* and *SCN10A,* important ion channels associated with pain signaling were significantly downregulated in the BoNT/A1 treated cells. (**Fig 4c**). Next, we examined Gene Ontology (GO) enrichment analysis for the top pathways selected based on the p-value (padj) ranking of the DEGs (**Fig 4e**). The enrichment analysis showed the top 3 downregulated biological pathways are associated with synaptic vesicle cycle, neurotransmitter transport, synaptic vesicle exocytosis (**Fig 4e**). Therefore, BoNT/A treatment on hDRG neurons, at least *in vitro*, has a large and unexpected effect on the transcription of a core set of genes that regulate neuronal transmission and excitability.

**Figure 4.**
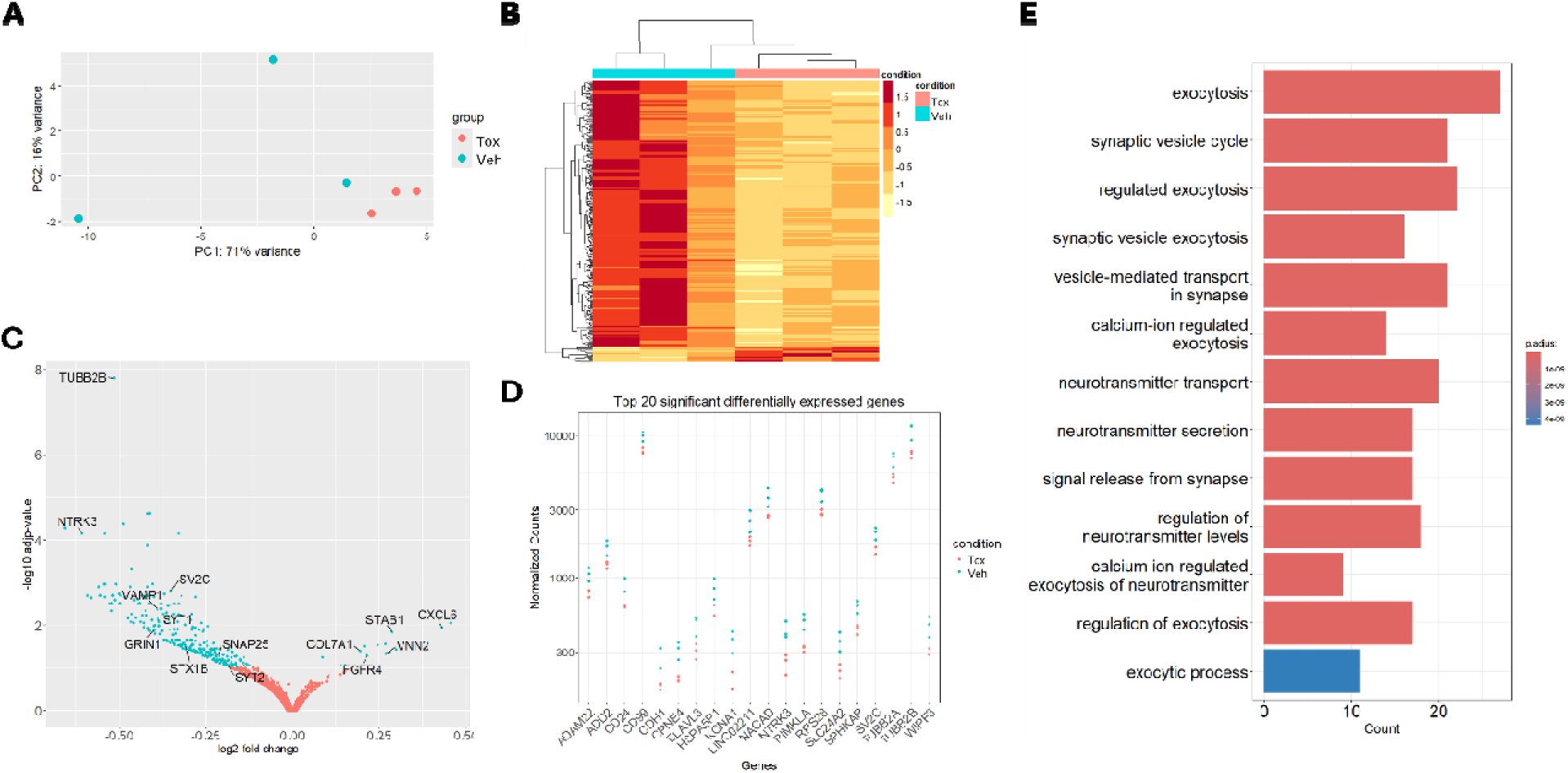
RNA sequencing on BoNT/A1 treated hDRG cultures. **A.** principal component analysis (PCA) plot showing the similarities and variability between the biological replicates and the two sample groups (Vehicle and Tox). **B.** Heatmap representing normalized values of *all* the significant genes between vehicle and toxin groups. **C.** Volcano plot illustrating the 273 differentially expressed genes between the vehicle and toxin groups. P values were adjusted for multiple comparisons. Labeled genes of interest with adjusted p < 0.01**. D.** plot displaying the normalized count values for the top 20 differentially expressed genes with adjusted p < 0.01. **E.** Representative GO enrichment analysis graph of DEGs; groups reflect main categories of GO term: BP, biological process.

### BoNT/A1 treatment decreases TRPV1-mediated CGRP release in hDRG neurons

To further investigate if BoNT/A1 treatment in hDRG neurons reduces neurosecretion like has been reported in rodent sensory neurons (Belinskaia *et al*. 2022; Edvinsson *et al*. 2015; Gazerani *et al*. 2010; Meng *et al*. 2009), we used an *in vitro* model of sensitized neurons treated with an inflammatory soup cocktail and treating them with the TRPV1 agonist capsaicin (Burstein *et al*. 1998; Sarchielli *et al*. 2006). The experimental timeline is shown in Fig 5a. We measured the CGRP secretion in the culture media via an ELISA. Compared to the untreated (veh/veh) cells, there was a significant decrease in CGRP release in the cells treated with BoNT/A1 (veh/toxin) (Fig 5b). In the group treated with inflammatory soup (soup/veh) we found an increase in CGRP release compared to the vehicle groups, demonstrating neuronal sensitization (Fig 5b). In the group treated with inflammatory soup and BoNT/A1 (soup/toxin), the CGRP release was significantly decreased by similar levels in the vehicle-toxin group (*** = p < 0.001; **= p < 0.01; by 1-Way ANOVA) (Fig 5b). To ensure that the inflammatory soup has not affected the cleavage of SNAP-25 in the toxin treated cells, we used immunocytochemistry to measure the cleavage of SNAP-25_197_, which was only detected in the BoNT/A1 treated groups in peripherin positive neurons (Fig. 5c**&d**). We conclude that BoNT/A1 decreases capsaicin-induced CGRP release both in naïve and sensitized hDRG neurons.

**Figure 5.**
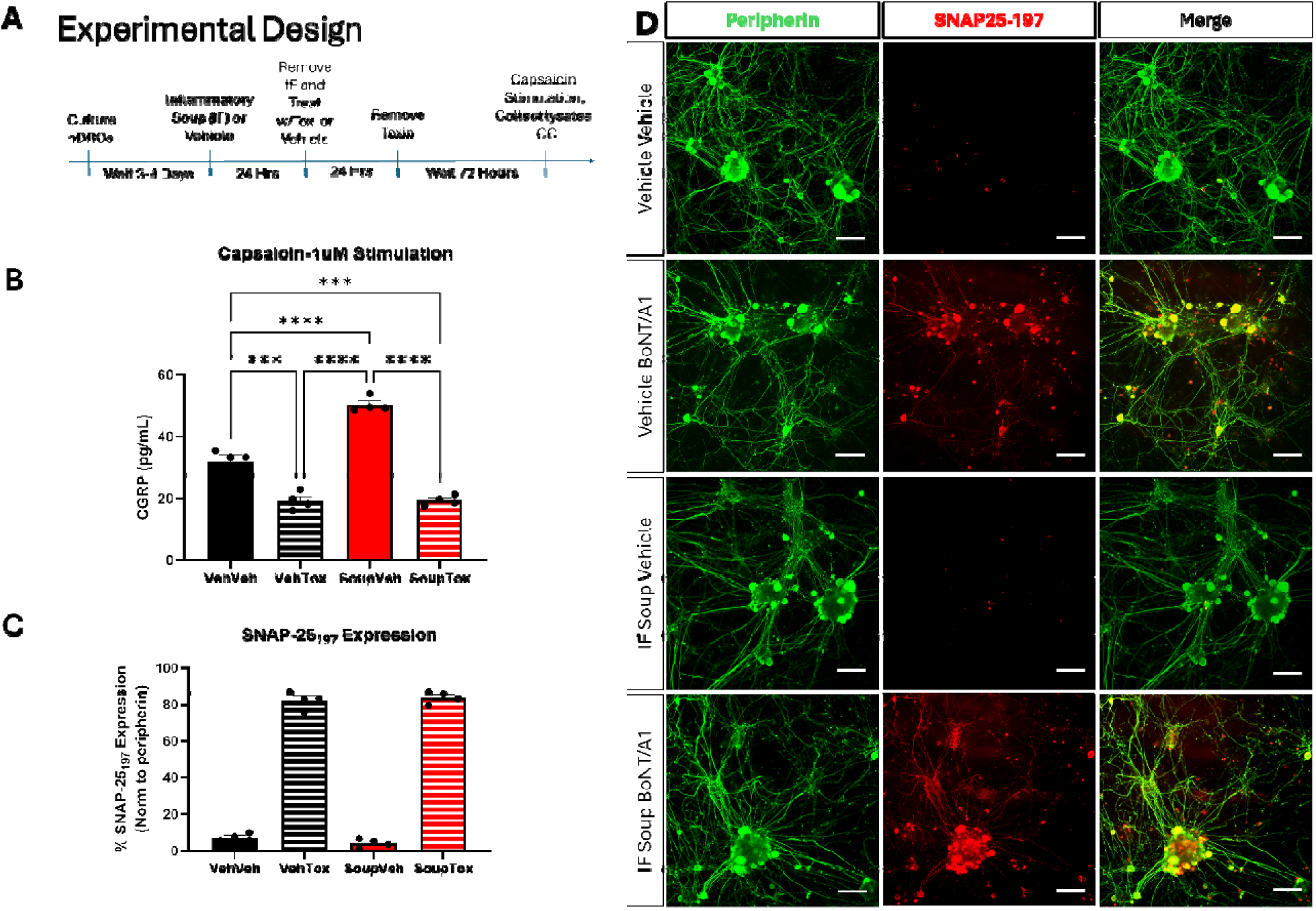
Capsaicin evoked CGRP release in hDRG neurons. **A.** experimental design outlining the inflammatory soup treatment or vehicle is given for 24 hours, followed by another 24-hour treatment of BoNT/A1 or vehicle. 72-hours later the hDRG neurons are treated with capsaicin (1uM) in Tyrodes media supplemented with 16uM thiorphan, a neutral endopeptidase (NEP) inhibitor, for30 minutes. The media was collected and CGRP release was measured by ELISA. **B**. Quantification of CGRP release (pg/mL) across all four groups (Veh/Veh, Veh/Tox, Soup/Veh, Soup/Tox) demonstrated that BoNT/A1 treatment (100pM) decreases CGRP release in naïve and sensitized neurons. *** = p < 0.001; **** = p < 0.0001; by 1-Way ANOVA with Tukey’s post-hoc test. **C.** Quantification of the % expression of cleaved SNAP-25_197_ normalized to peripherin positive neurons across all four treatment groups. Each data point represents the average of an individual replicate with at least 60 neurons counted across two different donors. Data is shown as means ± SEM. **D.** Representative 10x confocal images of previous quantification, showing peripherin, neuronal marker (Green), SNAP-25_197_ (Red), and their colocalization across all four treatment groups. The cleavage of SNAP-25 only occurs in the BoNT/A1treated neuronal cells. Scale bar=100um.

## Discussion

Our study provides new insights into the molecular mechanisms by which BoNT/A1 provides therapeutic benefits in human sensory neurons. Our findings align with previous rodent studies demonstrating that BoNT/A1 interferes with synaptic vesicle fusion and release of neurotransmitters and provide a robust examination of BoNT/A1’s effects on human nociceptors. Several key findings emerge from our experiments. The first is that contrary to previous work in hDRG tissues (Yiangou *et al*. 2011), we find that *SV2C* is the most highly expressed of the BoNT/A entry receptors in this tissue, far exceeding expression for *SV2A* or *SV2B*. Interestingly, the highest expression of *SV2C* was in Aβ nociceptors and silent nociceptors, two neuronal populations that play a key role in pain transmission (Djouhri & Lawson 2004; Lawson *et al*. 2019). Another key finding is the transcriptional change induced by BoNT/A1 in hDRG cultures. The discovery that BoNT/A1 treatment downregulates genes associated with neurotransmission and neuronal excitability suggests that some of the efficacy of BoNT/A in the clinic may be driven by this effect. Finally, we provide the first evidence that BoNT/A1 decreases baseline and evoked CGRP release in hDRG neurons. While this effect is expected given previous rodent studies (Belinskaia *et al*. 2022; Edvinsson *et al*. 2015; Meng *et al*. 2009), it supports a mechanism for BoNT/A1 to decrease vesicle mediated neuropeptide release in human nociceptive neurons.

Our findings confirm that BoNT/A1 cleaves SNAP-25 in a concentration-dependent manner in primary hDRG cultures, consistent with findings in rodent trigeminal ganglion neurons (Moore *et al*. 2023). SNAP-25_197_ was detectable in 80% of peripherin-positive neurons via immunocytochemistry and seen in cell lysates via Western blot confirming, by two different experimental approaches, that BoNT/A1 cleaves SNAP-25 in hDRG cells. Consequently, in our RNA sequencing experiments in cultured hDRG neurons, we identified 273 differentially expressed genes induced by BoNT/A1 treatment and GO enrichment analysis shows their involvement in synaptic vesicle cycling, neurotransmitter transport, and exocytosis.

Downregulation of key synaptic proteins such as *SNAP25, SV2C, SYT7, SYN2* and *VAMP* further suggests a transcriptional impact of BoNT/A1 on the molecular machinery required for synaptic vesicle fusion and neurotransmitter release. Neuronal markers such as CALCA, PIEZO2, NEFH, TRMP2 & F2RL2 remained unchanged as a response to BoNT/A1 treatment, further demonstrating specific regulation of synaptic protein trafficking and neurotransmitter release. While our work did not discover the mechanism of action responsible for this effect, it may provide insights into why multiple treatment cycles progressively reduce the headache days per month in some patients with chronic migraine treated with onabotA (Negro *et al*. 2015; Sarchielli *et al*. 2017) It may also be a clue into the therapeutic benefit of BoNT/A1 treatment for chronic migraine (Aurora *et al*. 2010; Aurora *et al*. 2011; Diener *et al*. 2010; Dodick *et al*. 2010; Lipton *et al*. 2011; Silberstein *et al*. 2013).

Among downregulated genes, we also found voltage-gated ion channels including *KCNA1* (Kv1.1), *CACNA1B* (Cav2.2) and *SCN10A* (NaV1.8) that may regulate neuronal excitability. These are crucial for action potential generation and are highly implicated in pain related disorders. Voltage gated potassium channel 1.1 (Kv1.1) has been found to act as a mechanical break to pain, and is a known target activated by the approved anti-inflammatory Niflumic acid (NFA) (Hao *et al*. 2013; Beekwilder *et al*. 2003; Servettini *et al*. 2023).

Downregulation of this channel would likely increase the excitability of sensory neurons and could be associated with a homeostatic mechanism to regulate the excitability of sensory neurons (McIlvried *et al*. 2025). The N-type voltage gated Calcium channel Cav2.2 is thought to be the major voltage gated calcium channel to regulate neurotransmitter release from nociceptors and the clinically approved drug, Ziconotide, targets this channel when given by intrathecal treatment of chronic pain (Winquist *et al*. 2005; Kutzsche *et al*. 2024; Brittain *et al*. 2011; Zamponi *et al*. 2015). Increased Nav1.8 channel expression can lead to spontaneous activity in sensory neurons and is thought to be a key mechanism of neuropathic pain (Gold *et al*. 2003; Roza *et al*. 2003; Lai *et al*. 2004). More recently, Suzetrigine a first in class Nav1.8 inhibitor was approved for treating acute pain (Jones *et al*. 2023; Vaelli *et al*. 2024; Osteen *et al*. 2025). BoNT/A1 downregulation of Nav1.8 and Cav2.2 would therefore be expected to have a robust effect on pain signaling by decreasing the expression of these crucial channels for pain. We acknowledge that future functional studies are needed to examine whether the downregulation of mRNA observed here leads to physiological changes in human nociceptors related to these ion channels.

The functional and translatability significance of these molecular alterations induced by treatment with BoNT/A1 was confirmed by measuring CGRP release from hDRG neurons. We chose to use a model that included sensitization of hDRG neurons by inflammatory soup followed by capsaicin stimulation and measured CGRP release, as CGRP is a well-established marker of nociceptive activation and transmission. BoNT/A1 treatment resulted in cleavage of SNAP-25 as well as a significant reduction of CGRP secretion in both naïve and sensitized hDRG neurons. This suggests that BoNT/A1 prevents neurotransmitter release by disrupting vesicle fusion, not only in non-sensitized neurons but also modulates the heightened neurotransmitter release observed in a sensitized state.

TRPV1 and P2X3 receptors have been shown to interact in a facilitatory manner to enhance pain signaling. Previous research has shown that BoNT/A treatment causes a reduction in transient receptor potential vanilloid 1 (TRPV1) and purinergic receptor P2X ligand-gated ion channel 3 (P2X3) in bladder of patients treated for overactive bladder (Coelho *et al*. 2010).

Given this, we examined our bulk RNA sequencing data for these two targets and found that *TRPV1* expression did not directly change in response to BoNT/A1 treatment transcriptionally. However, *P2RX3* was notably one of the downregulated genes in the toxin treated cells, suggesting a specific modulation of this receptor in sensory neurons following BoNT/A1 treatment. In the context of BoNT/A treatment, the reduction in P2X3 expression may disrupt its interaction with TRPV1, leading to an additional impact on TRPV1 cell surface expression or signaling (Saloman *et al*. 2013).

Overall, our results confirm that BoNT/A1 likely reduces the overall excitability of sensory neurons, preventing the excessive release of pain-related neurotransmitters. An alternative interpretation is that decreased TRPV1 function contributes to this effect, as discussed above and supported by findings from the only previous study that used hDRG neurons to study the effects of BoNT/A (Yiangou *et al*. 2011). However, our confirmation of the presence of SNAP-25_197_ in these experiments indicate that decreased vesicle fusion must contribute, at least partially, to the observed effects seen in our studies.

Collectively, our findings strongly suggest that BoNT/A likely enters human nociceptors through SV2C, where it cleaves SNAP-25 to disrupt synaptic vesicle trafficking, and subsequently ion channel expression and neurotransmitter release. A key finding from our work is evidence that treatment with BoNT/A causes transcriptional changes that presumably lead to downregulation of ion channels that play a crucial role in the transmission and modulation of pain signals and are important targets for the treatment of pain. The central strength of this study is the use of hDRGs; while TGs would be more directly relevant to migraines, their limited availability makes DRGs a valuable alternative for investigating BoNT/A’s general effects on sensory mechanisms. While our study provides additional foundational insights on BoNT/A’s impact on human sensory neurons, future research focusing on long-term treatment effects and impacts on neuronal plasticity in human tissues will be critical for optimizing its clinical application in pain management beyond migraine, such as neuropathic pain (Meyer-Friessem *et al*. 2019; Spagna & Attal 2023; Finnerup *et al*. 2015). This highlights the relevance of BoNT/A as a therapeutic agent in the human nervous system and its potential to modulate pain pathways. Future studies investigating whether Sv2c is essential for BoNT/A’s activity on human sensory neurons and its impact on CGRP release could deepen our understanding of the underlying mechanisms and enhance its therapeutic application in pain management.

## Author Contributions

PB-I., EV-C., designed and supervised research project and experiments. S.M., PB-I., conceptualized the manuscript. S.M., AB-A., LS., provided feedback and experimental input. KG and TJP wrote the manuscript, KH conducted the RNAscope, KG conducted the culturing and sequencing experiments. All authors edited the paper and provided input to the final manuscript.

## Data availability

All sequencing data will be deposited in GEO and made publicly available.

